# The oncogenic function of PLAGL2 is mediated via specific target genes through a Wnt-independent mechanism in colorectal cancer

**DOI:** 10.1101/2020.10.14.339531

**Authors:** Anthony D. Fischer, Daniel A. Veronese Paniagua, Shriya Swaminathan, Hajime Kashima, Deborah C. Rubin, Blair B. Madison

## Abstract

Colorectal cancer (CRC) tumorigenesis and progression are linked to common oncogenic mutations, especially in the tumor suppressor *APC*, whose loss triggers the deregulation of TCF4/β-Catenin activity. CRC tumorigenesis is also driven by multiple epimutational modifiers, such as transcriptional regulators. We describe the common (and near-universal) activation of the zinc finger transcription factor and Let-7 target PLAGL2 in CRC and find that it is a key driver of intestinal epithelial transformation. PLAGL2 drives proliferation, cell cycle progression, and anchorage-independent growth in CRC cell lines and non-transformed intestinal cells. Investigating effects of PLAGL2 on downstream pathways revealed very modest effects on canonical Wnt signaling. Alternatively, we find pronounced effects on the direct PLAGL2 target genes *IGF2,* a fetal growth factor, and *ASCL2*, an intestinal stem cell-specific bHLH transcription factor. Inactivation of PLAGL2 in CRC cell lines has pronounced effects on ASCL2 reporter activity. Furthermore, ASCL2 expression can partially rescue deficits of proliferation and cell cycle progression caused by depletion of PLAGL2 in CRC cell lines. Thus, the oncogenic effects of PLAGL2 appear to be mediated via core stem cell and onco-fetal pathways, with minimal effects on downstream Wnt signaling.

## INTRODUCTION

Colorectal cancer (CRC) is the third most common of all human malignancies and is the second leading cause of cancer-related deaths, after lung/bronchus cancer (seer.cancer.gov/statfacts). For over 20 years, the role of canonical Wnt signaling has been front and center for this malignancy, given the key role of the gatekeeper tumor suppressor, *APC*, which encodes a key scaffold protein in the β-Catenin destruction complex [1, 2]. Mutations in other Wnt pathway components, such as *AXIN2*, *CTNNB1*, and *RSPO* genes [3] can also lead to hyperactive Wnt signaling in CRC. Mutations in *TP53*, *SMAD4*, and *KRAS* are also very common in CRC, underscoring the roles of pathways regulating genome integrity, TGFβ/SMAD, and MAPK signaling. However, additional pathways are relevant in CRC pathogenesis, especially those pathways that drive an immature, fetal, or stem cell expression signature; such signatures predict an aggressive phenotype and poor prognosis [4, 5].

The Let-7 family of microRNAs (miRNAs) are well known for their key role in repressing naïve cellular states, controlling proliferation, and for maintaining cellular differentiation [6–9]. Consistent with this role Let-7 depletion promotes stem cell fate in intestinal epithelial cells [10] while down-regulation of Let-7 miRNA levels (or compromised Let-7 activity) fuels CRC carcinogenesis [10–12], with similar pro-oncogenic effects in many other malignancies [13–17]. Targets repressed by the Let-7 miRNA family are often part of proto-typical onco-fetal pathways that are frequently re-activated in a multitude of malignancies [8], including CRC [18, 19]. Such targets include *IGF2BP1*, *IGF2BP2*, *HMGA2*, *MYCN*, and a target we have recently characterized, *PLAGL2* [20]. We have previously documented the integral role of the Let-7 target HMGA2 in driving tumorigenesis in mouse models of intestine-specific Let-7 depletion [10], although HMGA2 alone does not appear to drive stem cell fate in intestinal epithelial cells [10]. In contrast, we discovered that PLAGL2 clearly drives stem cell fate in intestinal organoids, and directly activates the key stem cell-specific transcription factor, ASCL2 [20]. ASCL2 is critical for establishing and maintaining intestinal stem cell fate [21, 22].

Previous studies have demonstrated some oncogenic roles for PLAGL2 in CRC cell lines, with some documented effects on features of cellular transformation, *in vitro* [23–27]. Although these effects are hypothesized to be mediated via PLAGL2 enhancement of canonical Wnt signaling [23–27], the effects on Wnt signaling are often modest and the specific roles for such effects have not been determined. Studies in human CRC and glioma cells have revealed that PLAGL2 directly activates expression of *WNT6* [27, 28]. Despite this, effects on downstream Wnt signaling were not documented following manipulation of either PLAGL2 or WNT6 in CRC cell lines [27], whereas a clear effect was seen in neural stem cells [28]. This may reflect the commonplace ligand-independent hyperactivation of canonical Wnt signaling that occurs in CRC, e.g. from mutations in *APC*, which likely obscures (or renders irrelevant) any effects of individual upstream Wnt ligands. Further insight into the relationship between PLAGL2 and Wnt signaling was gained from experiments where we manipulated PLAGL2 levels in intestinal organoids [20]. PLAGL2 over-expression conferred Wnt-independent growth, but did not consistently augment Wnt signaling [20]. These results contrast the PLAGL2 studies described above, which hypothesize an obligatory downstream role for Wnt signaling in the context of cellular transformation. In non-transformed cells (organoids) we did find that factors (perhaps Wnt ligands) secreted from PLAGL2-overexpressing organoids enhanced a Wnt reporter in co-cultured organoids [20]. Consistent with this, PLAGL2 over-expression robustly activated expression of several Wnt ligands in organoids [20].

To investigate oncogenic mechanisms downstream of PLAGL2 we examined the potential of PLAGL2 to drive and maintain features of cellular transformation, and how such effects were mediated by direct PLAGL2 target genes. We find that PLAGL2 is a commonly up-regulated factor in CRC and has significant transforming properties. PLAGL2 effects on ASCL2- mediated signaling appear much more robust than effects on Wnt signaling. Through a close examination of TCF4/β-Catenin target genes and a TOP-tdT Wnt reporter, we find that PLAGL2 has minimal effects on canonical Wnt signaling. Consistent with these findings, over-expression of the PLAGL2 target genes *ASCL2* and *IGF2* (but not constitutive activation of Wnt signaling) can rescue proliferation defects caused by PLAGL2 inactivation.

## RESULTS

We first examined expression of *PLAGL2* in matched colonic adenocarcinomas to compare expression between tumors and non-malignant tissue from the same individual. *PLAGL2* expression was usually undetectable in non-tumor tissue (Fig 1A), whereas in tumor tissue *PLAGL2* was universally up-regulated (Fig 1A). Consistent with this, analysis of TCGA RNA-seq data for colorectal cancer revealed the common and robust up-regulation of *PLAGL2* in tumor tissue (Fig 1B). Expression analysis in CRC cell lines revealed *PLAGL2* expression in all CRC cell lines tested, with robust expression in HEK293T cells (Fig 1C). Further examination of TCGA data revealed that *PLAGL2* over-expression is a frequent feature of chromosomal instability (CIN) microsatellite-stable (MSS) tumors (P = 3.7e-17), a class that makes up the majority (∼80%) of CRCs. This is consistent with data from TCGA [29–31], where we find that the only mutations significantly associated with *PLAGL2* over-expression are *TP53* (P < 0.0001) and *APC* (P <0.01) (Fig 1D), which are hallmarks of CIN tumors [29]. This close association with *APC* loss could underlie observations that PLAGL2 expression correlates with β-Catenin levels in tumors [25]. As expected, *PLAGL2* up-regulation is rarely seen in tumors with mutations in *BRAF, ACVR2A, TGFBR2, MSH6,* or *MSH3* (P < 0.0001), which are typical in MSI tumors [29].

**Fig 1.**
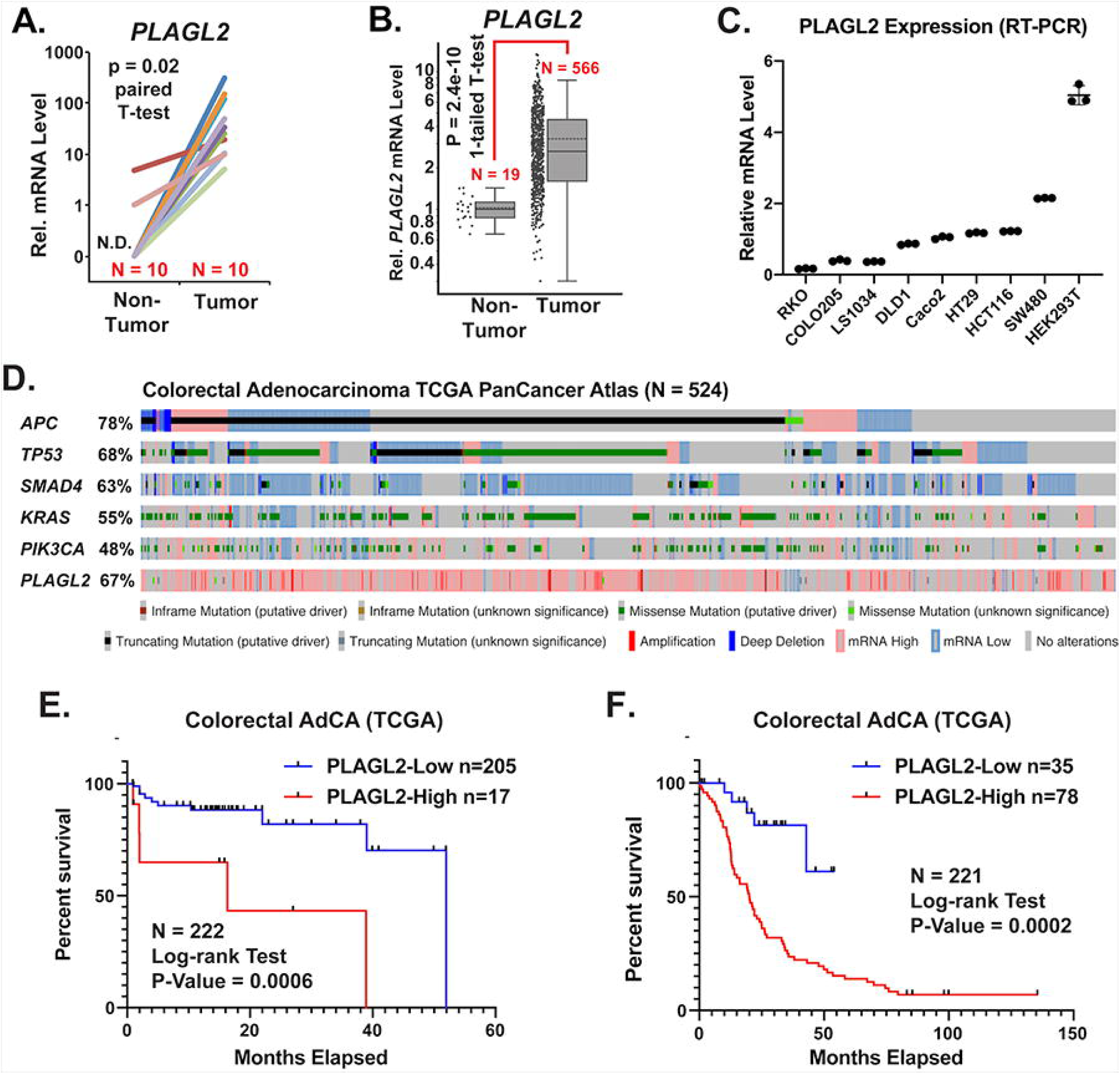
*PLAGL2* is turned on in the majority of colorectal cancers and is predictive of overall survival. **A)** *PLAGL2* mRNA expression in 10 pairs of colon adenocarcinomas, with matched non-tumor adjacent tissue, as previously described [10]. **B)** *PLAGL2* expression in non-tumor and colorectal adenocarcinoma tumors from the TCGA PanCancer Atlas cohort [69] **C)** RT-PCR assay quantifying relative levels of *PLAGL2* mRNA in colorectal cancer cell lines and HEK293T cells. **D)** Oncoprint showing mutation and expression profiles of commonly mutated cancer genes, including *PLAGL2*, from the TCGA PanCancer Atlas of colorectal adenocarcinomas. **E)** Overall survival of CRC patients stratified for high expression of PLAGL2 (>3 SD above mean) relativecompared to all other tumors, based on microarray expression data [29]. **F)** Overall survival of CRC patients stratified for high expression of PLAGL2 (>2 SD above mean) compared to low expression (>2 SD below mean) based on microarray expression data (TCGA Provisional). Statistical significance was evaluated using Student’s paired t-test **(A)**, Student’s one-tailed t-test **(B)**, or a Mantel-Cox log-rank test **(E, F)**.

To determine effects on disease outcomes, tumors were stratified for expression of PLAGL2 using data from two CRC TCGA cohorts, and analyzed for overall survival. In both cohorts, high expression of PLAGL2 in tumors predicted significantly reduced survival of CRC patients (Fig 1E, F). Thus, PLAGL2 is commonly turned on in CRC tumors, with high-level expression likely driving more aggressive disease progression.

We previously generated PLAGL2-mutant DLD1 cell lines [20] using SRIRACCHA [32] and here we examined their proliferation via a co-expressed H2BGFP reporter, which easily enabled their enumeration. Each mutant clone exhibited significant deficits in proliferation compared to non-mutant DLD1 parent cells (Fig 2A). While defined mutant clones generated with site-specific nucleases (such as CRISPR) can produce robust loss-of-function models, we also examined *PLAGL2* loss using CRISPR/Cas9 mutagenesis in a polyclonal population of mutant Caco2 and HT29 cells. Targeted cells were monitored using a transposon expressing a nuclear GFP reporter along with guide RNAs (gRNAs) against *PLAGL2* (Fig 2B). This transposon was delivered to cells already stably expressing EspCas9. In both Caco2 (Fig 2C) and HT29 (Fig 2D) cells, proliferation was significantly reduced in cells expressing the *PLAGL2*-specific gRNA. Lastly, shRNA knockdown of *PLAGL2* was pursued by a similar strategy with an shRNA-expressing transposon, along with a nuclear GFP reporter (Fig 2E). DLD1 cell lines established with this transposon revealed several shRNAs that robustly depleted *PLAGL2* mRNA (Fig 2F). In Caco2 cells expressing shRNA #3 and #4 (Fig 2G), proliferation was significantly reduced, especially for shRNA #4 (Fig 2H). Effects of *PLAGL2* knockdown were evaluated in the context of the FUCCI cell cycle reporter [33], delivered to cells using a transposon vector. In both Caco2 (Fig 2I) and HT29 (Fig 2J) cells, the FUCCI reporter revealed that *PLAGL2* knockdown decreased the number of cells in the S/G_2_/M phases of the cell cycle, but increased the number of cells in G_0_/G_1_. In sum, PLAGL2 depletion in CRC cell lines compromises proliferation and cell cycle progression.

**Fig 2.**
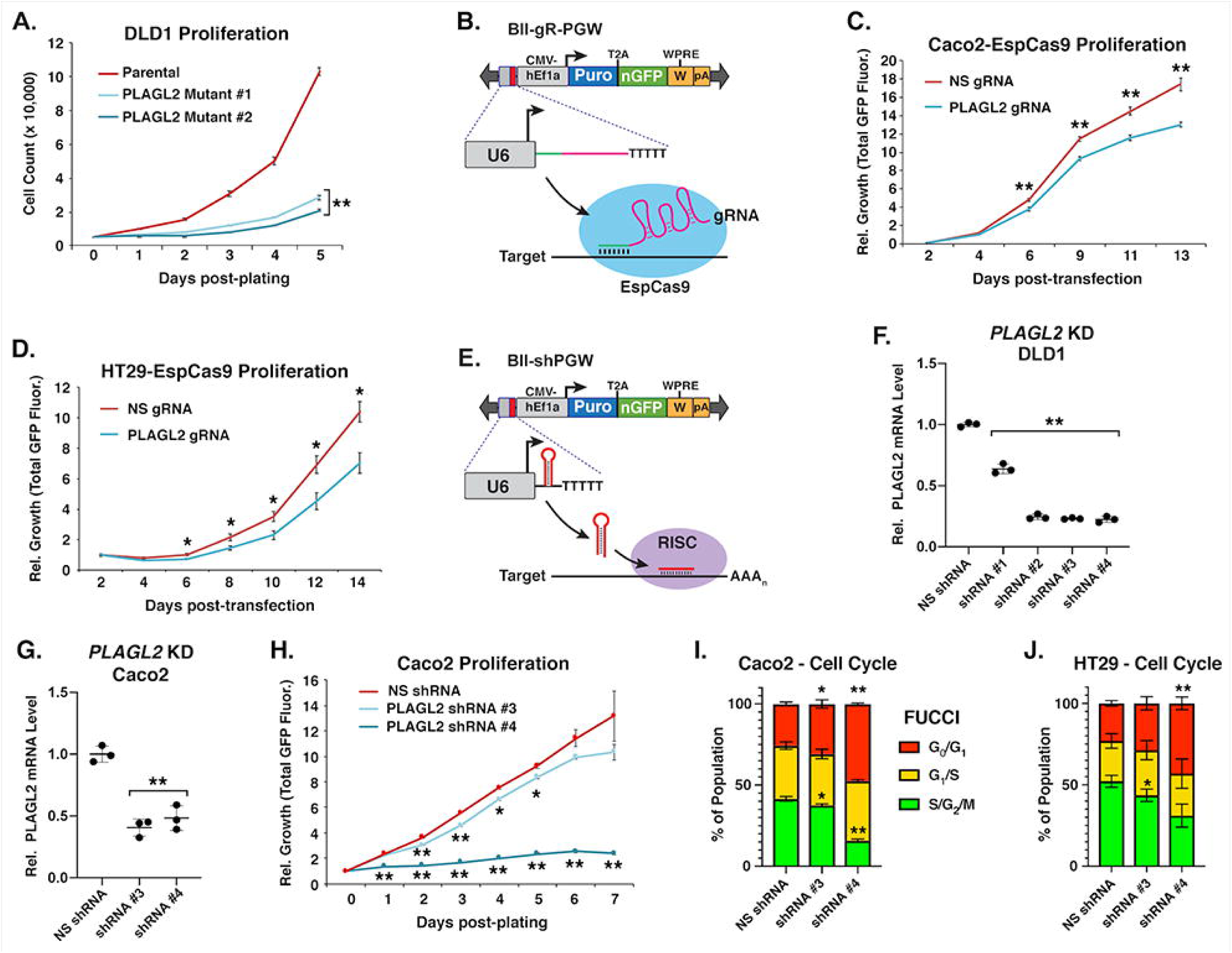
Knock-down or knock-out of *PLAGL2* compromises proliferation and cell cycle progression in CRC cell lines. **A)** Proliferation of stable CRISPR-mutated *PLAGL2* mutant DLD1 clones [20] compared to parental DLD1 cells. **B)** PB vector for stable expression of gRNAs and NLS-GFP, with Puro resistance. Caco2 **(C)** or HT29 **(D)** cell lines stably expressing EspCas9 were transfected with a PB vector expressing nuclear GFP and a non-specific (NS) gRNA or a gRNA against PLAGL2, and then growth was monitored. **E)** PB vector for stable expression of shRNAs and NLS-GFP, with Puro resistance. **F)** Knock-down (KD) of PLAGL2 was assessed by RT-PCR in DLD1 cells **(F)** and also confirmed in Caco2 **(G). H)** Proliferation of Caco2 cells stably transfected with PB vectors expressing nuclear GFP and non-specific (NS) or *PLAGL2*-specific shRNAs. Cell cycle status using the FUCCI [33] reporter co-transfected with a non-fluorescent shRNA PB vector was evaluated in Caco2 **(I)** and HT29 **(J)** CRC cells. For **A-D,** and **H** proliferation was quantified through enumeration of fluorescently labeled nuclei expressing NLS-GFP. Statistical significance (*p<0.05 or **p<0.01) was evaluated through an ordinary one-way ANOVA and Dunnett’s multiple comparisons post-hoc test.

Migration, invasion, and the ability to survive anchorage-independent growth are salient hallmarks of transformed cells. In transwell assays we evaluated migration in the DLD1 mutant clones and found that migration is severely compromised in both mutants (Fig 3A, D). Invasion through a layer of Matrigel was also remarkably reduced in DLD1 mutants, especially mutant #1 (Fig 3B, E, F). Knock-down of PLAGL2 in Caco2 cells also reduced migration in transwell assays, especially for shRNA #4 (Fig 3C). Lastly, *PLAGL2* was over-expressed in IEC6 cells to evaluate anchorage independent growth in soft agar. *PLAGL2* O/E consistently enabled the growth and formation of colonies (Fig 3G, H), demonstrating that PLAGL2 can confer resistance to anoikis-mediated cell death, a key property of transformed cells.

**Fig 3.**
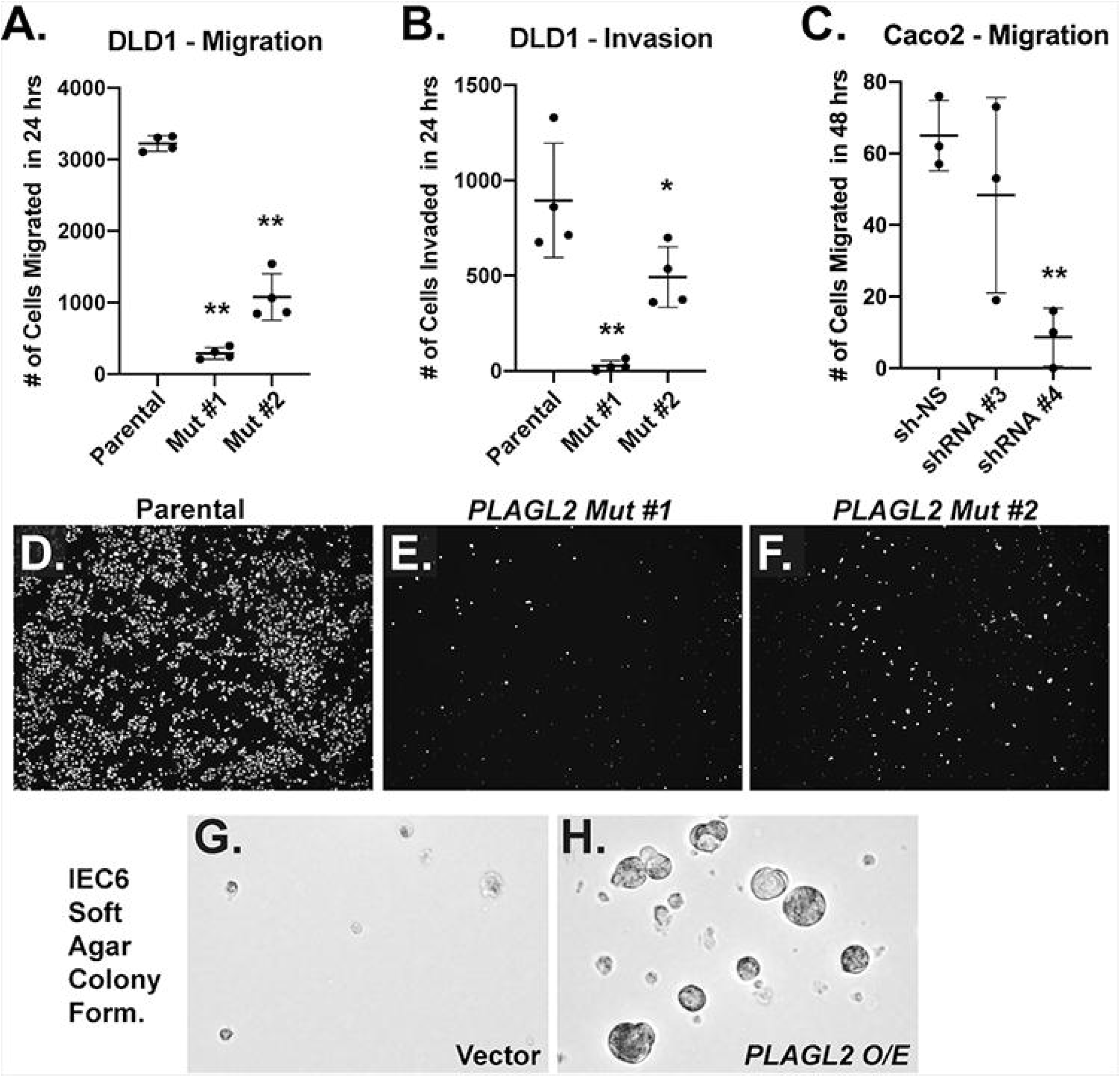
PLAGL2 drives migration, invasion, and colony formation in soft agar. Assays measuring migration **(A)** or invasion through Matrigel **(B)** of stable CRISPR-mutated *PLAGL2* mutant DLD1 clones [20] compared to parental DLD1. **C)** Migration assay of Caco2 cells stably transfected with PB vectors expressing nuclear GFP and a non-specific (NS) or a PLAGL2- specific shRNA. Images of GFP-expressing cells that have migrated through porous trans-well membranes 24 hours after plating for parental DLD1 cells **(D)**, DLD1 mutant clone #1 **(E)** or mutant clone #2 **(F)**. Soft agar colony forming assay for IEC6 cells transfected with a PB empty vector **(G)** or a PB vector expressing PLAGL2 **(H)**. Statistical significance (*p<0.05 or **p<0.01) was evaluated through an ordinary one-way ANOVA and Dunnett’s multiple comparisons post-hoc test.

We next investigated the role of specific PLAGL2 target genes. *IGF2* encodes a critical fetal growth factor that has been demonstrated to be a direct PLAG1 and PLAGL2 target gene [34–36], but its role downstream of PLAGL2 in the context of cellular transformation has not been investigated. Here we find that *IGF2* expression is significantly reduced in *PLAGL2* mutant DLD1 clones (Fig 4A) and following shRNA-mediated knockdown (Fig 4B). In SW480 cells we performed SRIRACCHA-enriched mutagenesis of *PLAGL2* with CRISPR/Cas9, delivered via transposon vector. RNA was extracted from a mixed polyclonal population of *PLAGL2* mutants for expression analysis, and mutagenesis of *PLAGL2* was confirmed by Illumina sequencing (S1 Fig). In this polyclonal population of *PLAGL2* mutants, *IGF2* expression was significantly reduced (Fig 4C). In defined clonal mutants, generated using CRISPR/Cas9 (also delivered via transposon vector) in mouse organoids [20] *Igf2* expression was also significantly reduced. Thus, *PLAGL2* is required for expression of *IGF2* in both transformed and non-transformed intestinal epithelial cells. Over-expression of PLAGL2 in mouse organoids [20] resulted in robust dose-dependent up-regulation of *Igf2* mRNA (Fig 4E). In sum, PLAGL2 appears necessary and sufficient to drive *IGF2* expression. To gauge effects of IGF2 alone, the human IGF2 cDNA was over-expressed in mouse intestinal enteroids via transposon transgenesis (Fig 4F). IGF2 expression triggered organoid hyperplasia and the formation of large cysts (Fig 4G, H), which phenocopies the cyst-like appearance of PLAGL2-overexpressing enteroids [20].

**Fig 4.**
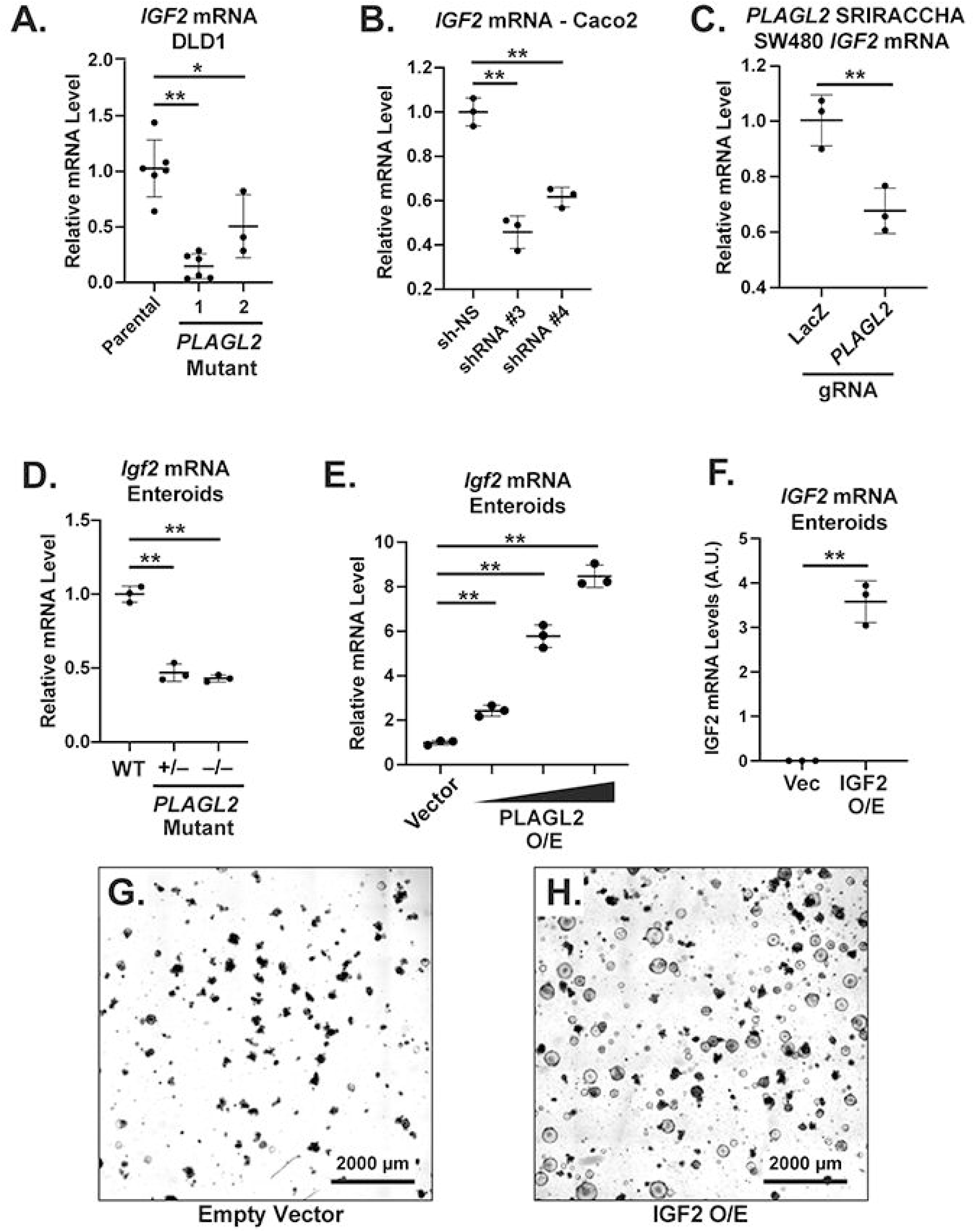
PLAGL2 drives IGF2 expression in CRC cells and intestinal organoids and partially rescues growth phenotype in *PLAGL2* mutant CRC cells. **A)** RT-PCR for *IGF2* in DLD1 *PLAGL2* mutant clones. **B)** RT-PCR for *IGF2* in Caco2 cells following shRNA KD of *PLAGL2*. **C)** RT-PCR for *IGF2* following CRISPR/Cas mutagenesis using SRIRACCHA [32] and Hygromycin selection. **D)** RT-PCR for *Igf2* in *Plagl2* knockout mouse intestinal enteroids [20] compared to WT parental enteroids. **E)** RT-PCR for *Igf2* in 3 lines of mouse intestinal enteroids expressing low, medium, or high levels of human HA-tagged PLAGL2 [20] compared to empty vector control enteroids. **F)** RT-PCR for IGF2 mRNA following O/E in enteroids. **G+H)** Cyst-like morphology in IGF2 O/E enteroids. For comparison of more than 2 conditions the statistical significance (*p<0.05 or **p<0.01) was evaluated through an ordinary one-way ANOVA and Dunnett’s multiple comparisons post-hoc test. For pair-wise comparison, Welch’s t-test was performed to determine statistical significance (*p<0.05 or **p<0.01).

To investigate functional roles of PLAGL2 target genes, the downstream effectors *ASCL2* and *IGF2* were further examined in tumors and transformed CRC cell lines. We examined CRC tumors from the TCGA for expression correlation between *PLAGL2* and ASCL2 target genes (*ASCL2, KLHDC4, OLFM4, RNF43, LGR5, MYB, NR2E3, SMOC2, OSBPL5,* and *SOX9*). Nine out of ten ASCL2 targets showed positive correlation with *PLAGL2* in all three TCGA CRC cohorts, while most targets were also positively correlated with *PLAGL2* in stomach adenocarcinoma and hepatocellular carcinoma tumors (Fig 5A). Consistent with this, CRC cell lines stratified for the lowest vs. highest quintile of PLAGL2 expression, as determined by previous RNA-seq studies [37], revealed that expression of *ASCL2* is significantly higher in *PLAGL2*-high CRC cell lines (Fig 5B). SRIRACCHA-enriched mutagenesis of *PLAGL2* in SW480 cells (S1 Fig) also revealed depletion of ASCL2 mRNA levels. This is all consistent with a role for PLAGL2 in the direct transcriptional activation of *ASCL2*, as previously demonstrated in intestinal organoids and CRC cell lines [20]. However, to quantitatively gauge the impact on ASCL2 activity in CRC cell lines we used the ASCL2 reporter [38], adapted for transposon-mediated expression of tdTomato (STAR-tdT, Fig 5D). Co-transfection with shRNA vectors or an ASCL2 vector revealed robust activation by ASCL2 and modest reduction by shRNAs against *PLAGL2* (Fig 5E). After establishing stable transgenic lines, shRNA knockdown of *PLAGL2* caused a pronounced reduction in STAR-tdT reporter activity in SW480, HT29, and Caco2 cells (Fig 5F-H).

**Figure 5.**
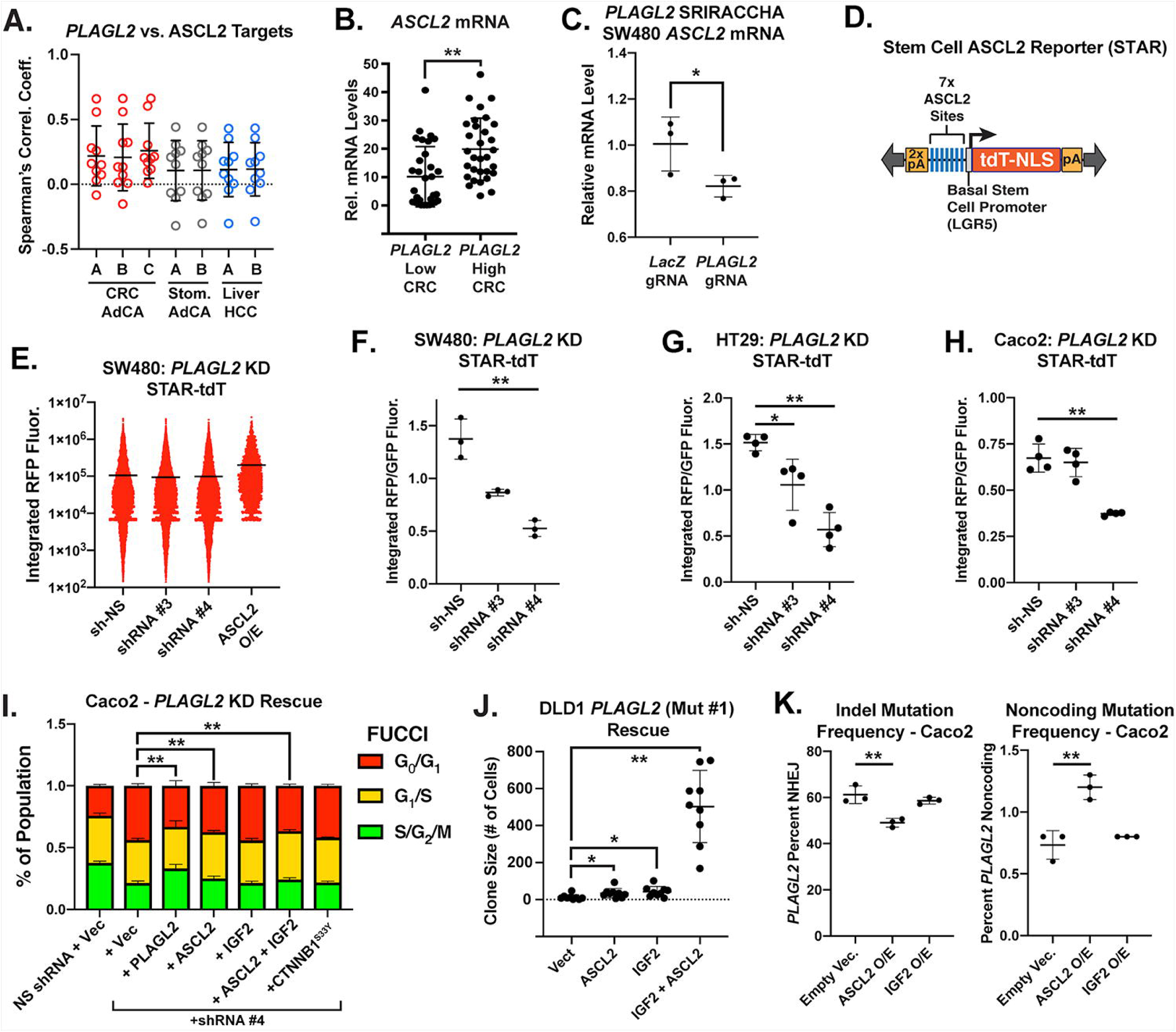
PLAGL2 activates an ASCL2-dependent program in CRC cells. **A)** Plots of correlation coefficients of linear regressions between *PLAGL2* and ten ASCL2 direct target genes (*ASCL2, KLHDC4, OLFM4, RNF43, LGR5, MYB, NR2E3, SMOC2, OSBPL5,* and *SOX9*) for seven TCGA datasets, including 3 from CRC adenocarcinoma (AdCA), 2 from stomach AdCA, and 2 from liver hepatocellular carcinoma (HCC). **B)** Colorectal cancer cell lines (N=154 from [37]) were parsed into bottom and top quintiles for *PLAGL2* expression and then evaluated for *ASCL2* mRNA expression. **C)** EspCas9-mediated mutagenesis using SRIRACCHA [32] and gRNAs directed against LacZ or *PLAGL2*, followed by selection with hygromycin and RT-PCR for *ASCL2* mRNA. **D)** Depiction of the STAR ASCL2 reporter construct. **E)** Activity of the stem cell ASCL2 reporter (STAR-tdT) reporter [38] 48 hours after transfection in SW480 CRC cells that were also co-transfected along with vectors providing *PLAGL2* shRNA-mediated KD or ASCL2 O/E. As expected ASCL2 O/E augments STAR-tdT reporter activity. After selection with Puromycin for 6-8 days for stable *PLAGL2* shRNA expression STAR-tdT reporter activity (integral RFP fluorescence) was measured in SW480 **(F)**, HT29 **(G)**, and Caco2 **(H)** CRC cell lines. RFP levels were normalized to integral GFP fluorescence constitutively expressed by the PB shRNA vector. **I) FUCCI** analysis of cell cycling in Caco2 cells after shRNA mediated KD of *PLAGL2* and exogenous expression of PLAGL2, ASCL2, IGF2, ASCL2 and IGF2, or a constitutively active β-catenin. **J)** Rescue of DLD1 *PLAGL2* mutant clone #1 with IGF2 and/or ASCL2 O/E and assessment of colony size following selection with G418. **K)** Mutation profile of PLAGL2 targeted with SRIRACCHA using eSpCas9 with simultaneous over-expression of ASCL2 or IGF2. For comparison of more than 2 conditions the statistical significance (*p<0.05 or **p<0.01) was evaluated through an ordinary one-way ANOVA and Dunnett’s multiple comparisons post-hoc test. For pair-wise comparison, Welch’s t-test was performed to determine statistical significance (*p<0.05 or **p<0.01).

If ASCL2 and/or IGF2 are critical drivers downstream of PLAGL2, then we expect that their expression would rescue growth following PLAGL2 loss-of-function. We first examined proliferation using the FUCCI reporter following knock-down of PLAGL2 and rescue with ASCL2 and/or IGF2. Only ASCL2 was able to restore cell cycle progression in Caco2 cells while neither IGF2 nor constitutively active β-Catenin (*CTNNB1^S33Y^*) expression was sufficient (Fig 5I). Because of their expression of H2BGFP, DLD1 *PLAGL2* mutants could not be assessed using the FUCCI reporter. However, in *PLAGL2* DLD1 mutant line #1, ASCL2 and IGF2 individually augmented clone size following stable transfection, and when co-transfected together we observed synergistic effects between ASCL2 and IGF2 on growth in this *PLAGL2* mutant (Fig 5J), suggesting a role for both target genes. To examine a more diverse array of *PLAGL2* mutants, we performed SRIRACCHA-mediated mutagenesis [32] in Caco2 CRC cell lines while simultaneously providing PB-mediated transgenic expression of either IGF2 or ASCL2. If either IGF2 or ASCL2 is able to compensate for loss of PLAGL2, then we would expect increased survival and growth of clones with *PLAGL2* loss-of-function mutations if such clones also express exogenous IGF2 or ASCL2. Thus, a rescue would be evident in a higher proportion of *PLAGL2* mutants. Exogenous IGF2 expression does not result in a higher frequency of *PLAGL2* mutations, but surprisingly, ASCL2 expression results in a significantly lower frequency of *PLAGL2* mutations and higher proportion of non-coding mutations (Fig 5K). In a 2-step experiment where lines were first established that already over-expressed ASCL2 or IGF2, *PLAGL2* mutagenesis was not tolerated in Caco2 cells already over-expressing ASCL2; i.e. no clones were recovered, unlike IGF2 over-expressing or vector controls. This suggests that any oncogenic effects of ASCL2 depend on PLAGL2, and that these two factors may function as obligate partners.

Previous studies have reported that PLAGL2 augments Wnt signaling [23, 24, 27], perhaps via the transcriptional activation of Wnt ligand genes [28]. To explore effects of PLAGL2 on canonical Wnt signaling, we compared expression levels of *PLAGL2* mRNA with 36 Wnt target genes among 7 cancer datasets from the TCGA (Fig 6A). While some targets showed a positive correlation, many targets also showed anti-correlation with PLAGL2, and we observed no overall trend towards a positive association, except perhaps a slight positive correlation among liver hepatocellular carcinoma samples (Fig 6B). In addition, an examination of fold changes of these Wnt targets in RNA-seq data from PLAGL2-expressing intestinal organoids showed no clear trend (Fig 6C). In contrast, data sets directly manipulating Wnt signaling showed a clear decrease in Wnt target expression following β-Catenin knock-down or dnTCF4 expression, while stimulation of Wnt signaling in intestinal organoids (via GSK3β inhibition) showed an induction of Wnt target gene expression (Fig 6C). Canonical Wnt signaling is also frequently measured via heterologous reporters. We used our TOP-tdT reporter [20] to quantify Wnt signaling in the context of PLAGL2 knock-down. Co-transfection of dnTCF4 dramatically reduced signal from this reporter, as expected (Fig 6D). After establishing stable shRNA expression in the context of the TOP-tdT reporter, we observe a small decrease in normalized reporter signal in SW480 and Caco2 cells (Fig 6E, F), but not in HT29 cells (Fig 6G). Representative images from Caco2 experiments confirm modest effects on TOP-tdT signal (Fig 6H). Despite subtle effects in SW480 and Caco2 cells, no significant effects were observed on non-phosphorylated (“active”) β-Catenin or total β-Catenin protein levels in these cells (Fig 6I, J). In DLD1 *PLAGL2* mutant cell lines, similar effects were observed. Values were quantified from these immunoblot experiments (Fig 6 K-M). Expression analysis of Wnt target genes also did not reveal any clear trend following *PLAGL2* knock-down (Fig 6N-P). SRIRACCHA-mediated mutagenesis of *PLAGL2* in SW480 cells (S1 Fig) and RT-PCR revealed a slight trend of reduced Wnt target gene expression, but only *ETS2* mRNA levels were significantly reduced (Fig 6Q, R). In sum, the data only support a small effect of *PLAGL2* on Wnt signaling in CRC cells.

**Fig 6.**
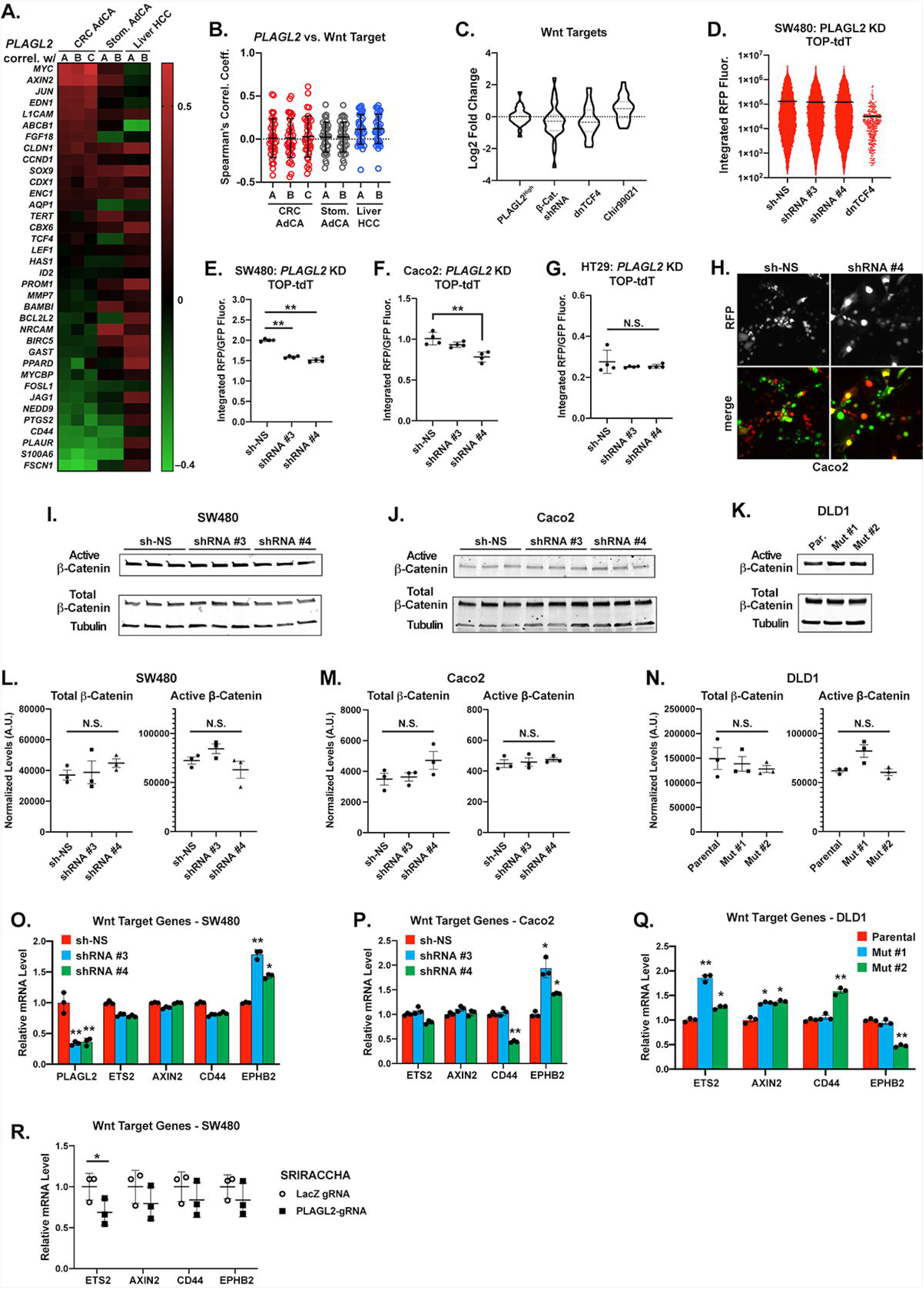
Minimal effects on canonical Wnt signaling following inactivation of PLAGL2. **A)** Heatmap of correlation coefficients for linear regressions between *PLAGL2* and 36 TCF4/β-Catenin target genes for seven TCGA datasets, including 3 from CRC adenocarcinoma (AdCA), 2 from stomach AdCA, and 2 from liver hepatocellular carcinoma (HCC). **B)** Dot/scatter plots of data from **(A). C)** Violin plots of fold changes among 28 TCF4/β-Catenin target genes for conditions as follows: 1) between PLAGL2-low vs. PLAGL2-high CRC AdCA tumors (TCGA), 2) between control and KD of *CTNNB1* (β-Catenin) or inducible O/E of a dnTCF4 in LS174T CRC cell lines [70], and 3) GSK3β inhibition and Wnt/TCF4/β-Catenin activation in intestinal organoids following treatment with 5 µM CHIR99021 [71]. **D)** Activity of the Wnt Super 8x TopFlash reporter [72] driving expression of tdT (TOP-tdT) 48 hours after transfection in SW480 CRC cells that were also co-transfected along with vectors providing *PLAGL2* shRNA-mediated KD or dnTCF4 O/E. As expected, dnTCF4 O/E reduces signal from the TOP-tdT reporter and reduces the number of RFP-positive cells. After selection with Puromycin for 6-8 days for stable *PLAGL2* shRNA expression TOP-tdT reporter activity (integral RFP fluorescence) was measured in SW480 **(E)**, Caco2 **(F)**, and HT29 **(G)** CRC cell lines. RFP levels were normalized to integral GFP fluorescence constitutively expressed by the PB shRNA vector. **H)** Representative images of fluorescent Caco2 cells after 6 days of selection with 10 µg/mL puromycin. Active (non-phosphorylated) and total β-Catenin levels were measured following *PLAGL2* KD in SW480 **(I)**, Caco2 **(J)**, and *PLAGL2* mutant DLD1 clones **(K)**. Protein levels were quantified and normalized to tubulin **(L-N)**. Direct TCF4/β- Catenin target gene expression levels were measured by RT-PCR in SW480 and Caco2 cells following PLAGL2 KD **(O, P)**, in *PLAGL2* mutant DLD1 clones **(Q)**, and following SRIRACCHA mediated mutagenesis of *PLAGL2* in SW480 cells **(R)**. Statistical significance (*p<0.05 or **p<0.01) was evaluated through an ordinary one-way ANOVA and Dunnett’s multiple comparisons post-hoc test.

## DISCUSSION

We find that PLAGL2 is a transcriptional regulator of cellular transformation in colorectal cancer, being able to drive hallmark features, including unchecked proliferation, migration, invasion, and anchorage independent growth. The normal physiological role of PLAGL2 is likely integral to a developmental pathway considering its high expression in the fetal intestinal anlagen and early postnatal intestine, but low expression in adult tissue [39]. *Plagl2*^−/−^ mice demonstrate defects in intestinal epithelial differentiation and function, suggesting that PLAGL2 plays a critical role during fetal intestinal development [39]. Similar to many other Let-7 targets, such as *HMGA2*, *PLAGL2* appears to exhibit typical features of an onco-fetal gene, considering its frequent re-activation in CRC and other malignancies [20, 28, 40–42]. However, there has been little insight into the role of direct transcriptional targets of PLAGL2, either during fetal development or carcinogenesis. Our current studies here were all conducted *in vitro*, which have many limitations, and validating *in vivo* models should be pursued in the future to delineate the role of PLAGL2 in intestinal mucosal transformation.

In possible oncogenic roles, PLAGL2 has been shown to directly activate transcription of the thrombopoietin receptor (*MPL*) [43], which is a key receptor necessary for megakaryocyte and platelet formation [44]. MPL is also a proto-oncogene that can drive hematopoietic cell proliferation when it (or the truncated *v-mpl* oncogene) is over-expressed [45, 46]. Relevant to PLAGL2, MPL is implicated in driving transformation downstream of PLAGL2 in acute myeloid leukemia (AML) [43], a malignancy in which increased expression of MPL marks a particularly aggressive subset [47]. However, even though MPL has been shown to be expressed on a subset of CRC cells that have metastatic tropism for the liver and lung [48, 49], *MPL* expression is not induced by PLAGL2 in intestinal organoids [20] and *MPL* expression does not correlate with *PLAGL2* expression in CRC [29]. Thus, *MPL* may not be a relevant oncogenic PLAGL2 target gene in CRC. Other potential PLAGL2 targets, such as *NIP3* and *P73* [50, 51], appear to have tumor-suppressive properties, while another documented target, surfactant protein C (*SPC*) [52], has unknown relevance to tumorigenesis and/or cellular transformation.

We have previously identified *ASCL2* as a direct target gene of PLAGL2. ASCL2 is a bHLH transcription factor that directly supports *LGR5* expression [22], drives IESC fate [21, 22, 53], and promotes an aggressive phenotype in CRC [53–55]. Regarding this effect in CRC, ASCL2 represses expression of *CDX2*, a transcription factor with a well-described role in positively driving intestinal epithelial differentiation [54]. ASCL2 also represses expression of miR-200 miRNAs, which may underlie effects on a mesenchymal phenotype and/or EMT in cancer, which is repressed by the miR-200 family of miRNAs [55]. Studies have also suggested that ASCL2 may play a role in augmenting the tumor-initiating capacity of CRC cells [56, 57], although this has not yet been carefully examined. Ultimately, the role of *ASCL2* may prove to be pleiotropic, as these studies suggest, with variable effects on differentiation, EMT, and tumor initiating potential.

Surprisingly, in our studies, ASCL2 over-expression confers a survival disadvantage to *PLAGL2* mutants in the Caco2 cell line, even though we see that cell cycle progression is promoted by ASCL2 rescue in the context of PLAGL2 knock-down. Thus, ASCL2 alone may have different effects than when co-expressed with PLAGL2 at high levels; such effects of ASCL2 may repress cellular transformation when PLAGL2 levels are low. Previous studies seeking to identify ASCL2 co-immunoprecipitating proteins included the identification of PLAGL2, as determined by mass spectrometry [38], although such direct interaction was not verified by other methods. If direct interaction can occur between these factors, then the effects we see on the STAR-tdT reporter in CRC cell lines could possibly reflect direct roles for PLAGL2 on ASCL2-responsive regulatory elements. While follow-up studies are needed, PLAGL2 and ASCL2 may biochemically cooperate to drive specific oncogenic targets – targets that may differ from those that are transcriptionally activated individually by each individual factor.

For effects on tumorigenesis, a gain-of-function role for ASCL2 (e.g. in the context of *ASCL2* up-regulation) is not clear. Modest entopic over-expression of *Ascl2*, at levels 2-3-fold higher than non-transgenic mice, via a BAC transgene, does not accelerate CRC tumorigenesis in the context of the *Apc^Min^* allele [58], a tumor model in which mice develop adenomatous polyps in the small intestine and colon. However, DNA replication (as measured by bromodeoxyuridine incorporation) is significantly elevated in the intestinal epithelium of these *Ascl2* transgenic mice, suggesting that elevated levels of ASCL2 can drive cell cycle progression [58]. Consistent with these *in vivo* findings, we observe accelerated cell cycle progression following ASCL2 over-expression in the context of PLAGL2 knock-down. In contrast to SW480 cells, which express very low levels of ASCL2 mRNA and protein, Caco2 cells express very high levels of ASCL2 [54]. Thus, ASCL2 may play a differential role downstream of PLAGL2, depending on the individual tumor or cancer cell line. In light of *in vivo* studies described above, *ASCL2* may only have cancer-promoting effects when *PLAGL2* is also coordinately up-regulated and/or activated. The co-expression of *PLAGL2* with *ASCL2* is striking in CRC [20], so the context for cooperation certainly exists in tumors.

The PLAGL2 paralog, PLAG1, is reported to transcriptionally activate the IGF2 promoter [34, 36], with over-expression experiments in NIH-3T3 and HEK293 cells suggesting that PLAGL2 may also positively regulate *IGF2* expression. IGF2 (a known driver of tumor progression) is over-expressed in ∼15% of CRCs and also activates the PI3K-AKT pathway [59–65]. IGF2 up-regulation also occurs in the context of wild-type *PTEN*, *PIK3CA*, *BRAF*, and *KRAS* [29], suggesting that IGF2 over-expression can functionally substitute for common PI3K-AKT-activating mutations. Here we provide data in primary intestinal organoids and human CRC cell lines that PLAGL2 is necessary and sufficient to drive *IGF2* expression. Following knock-out or knock-down of *PLAGL2* in CRC cell lines, we observe that *IGF2* expression is depleted up to 95%, and in a CRC cell line such as Caco2, which is known to express very high levels of IGF2 [66–68]), severe reduction of IGF2 has consequential effects on cellular growth. Evidence for this exists through studies of Caco2 cells, which are sensitive to both a neutralizing anti-IGF2 antibody [66] and an IGF1R/INSR inhibitor [68], which blocks IGF2 or the receptors through which IGF2 signals, respectively. However, we cannot rescue cellular proliferation and/or survival following *PLAGL2* mutagenesis in Caco2 cells via over-expression of IGF2. Thus, the growth defects caused by PLAGL2 loss are due to other effectors, besides IGF2. In *PLAGL2* mutant DLD1 cells ASCL2 and IGF2 each alone have modest effects on clone proliferation, but cooperative/synergistic effects when co-expressed. Therefore, in some contexts these PLAGL2 targets cooperate, whereas in other contexts, such as Caco2 cells, we do not see evidence of cooperation. Thus, the oncogenic dependency of PLAGL2 on IGF2 and ASCL2 is context-dependent, varying between tumors and/or tumor types.

Finally, our data suggest a minor role for Wnt signaling downstream of PLAGL2 in CRC. While previous studies indicate that WNT6 is a PLAGL2 target gene [27, 28], and our previous studies show that PLAGL2 activates the expression of Wnt genes (i.e. *Wnt9b*, *Wnt4*, *Wnt10a*, and *Wnt5a*) in primary intestinal epithelial cells [20], our studies here in CRC cell lines indicate that canonical Wnt signaling (via TCF4/β-Catenin) are only modestly affected by PLAGL2. Additionally, in available data from TCGA we find that PLAGL2 levels do not correlate with a Wnt signature, as measured by the induction of Wnt target genes. If the effects of PLAGL2 on Wnt signaling are routed via Wnt ligands (such as WNT6), then a minor effect on Wnt activation is not unexpected, since the vast majority of colorectal cancers possess mutations in either *APC*, *AXIN2*, or *CTNNB1* [29] — mutations that lead to ligand-independent Wnt pathway activation. However, an earlier role may exist for Wnt-dependent effects of PLAGL2 if PLAGL2 is activated prior to cell autonomous activation of intrinsic Wnt signaling through such mutations. This is a limitation of our study – the inability to model early pre-cancerous lesions prior such canonical Wnt pathway mutations. Other PLAGL2 target genes, besides Wnt ligands themselves, could be intracellular modifiers of the canonical Wnt signaling pathway. However, given the modest effects of PLAGL2 on canonical Wnt signaling in CRC cells, the net effect of any such hypothetical targets is likely to be relatively small. This underscores the need to identify additional PLAGL2 target genes, especially those involved in fetal growth pathways and carcinogenesis; identifying such targets will help illuminate the onco-fetal pathways downstream of PLAGL2.

## MATERIALS AND METHODS

### Statistical Analysis

All statistical analyses were performed using GraphPad Prism (Version 8 for Windows, GraphPad Software, San Diego, California USA, www.graphpad.com). Threshold for significance (alpha level) was 0.05 for all tests.

### Cell lines

Cell lines used in this study include Caco-2 [Caco2] (ATCC® HTB-37™), HT29 [HT-29] (ATCC® HTB-38™), SW480 [SW-480] (ATCC® CCL-228™), DLD1 [DLD-1] (ATCC® CCL-221™), and IEC6 [IEC-6] (ATCC® CRL-1592™). Cells were cultured in media prescribed by ATCC with the inclusion of a prophylactic dose of Plasmocin (Invivogen) to prevent mycoplasma contamination.

### Ethics statements

Colorectal tumor specimens were provided by the Siteman Cancer Center Tissue Procurement Core, Human Studies Institutional Review Board number 201106191. These tissues were fully anonymized prior to accessing them, thus an individual IRB was not required.

Mice are housed in the fully AAALAC/OLAW certified animal facility at the Washington University in St. Louis School of Medicine. Mice are housed in a 12 hour light/dark cycle with free access to food and water; the number of mice per cage was as mandated by the Institutional Animal Care and Use Committee (IACUC) of Washington University in St. Louis School of Medicine to ensure humane caging. All animal experimentation was approved by the IACUC/ Animal Studies Committee of Washington University School of Medicine as stated in approved protocol # IACUC 20170164.

Euthanasia was performed by C02 asphyxiation with a flow control gauge followed by cervical dislocation as recommended and approved for mice by the American Veterinary Medical Association Panel on Euthanasia and by the IACUC and Division of Comparative Medicine at Washington University in St. Louis School of Medicine.

### RT-PCR

CRC tumor samples, along with paired non-cancerous controls, were obtained from Siteman Cancer Center Tissue Procurement Core and were previously described [10]. RNA was extracted with TRIzol (ThermoFisher Scientific) and further purified using the RNeasy RNA Clean-up Kit (Qiagen) for these tumor samples. For cell lines or enteroids, RNA was prepared using TRIzol. RT reactions were performed with 3-4 µg total RNA using oligo-dT and SuperScript III RT (ThermoFisher Scientific). QPCR was performed as previously described [20], and expression levels normalized to *PPIA* and *B2M* for human specimens or human cell lines. For RT-PCR using RNA from mouse organoids or cell lines expression levels were normalized to *Hprt* and *Tbp*.

### TCGA analysis

TCGA data was examined using the cBioPortal for Cancer Genomics [30, 31], hosted by the Center for Molecular Oncology at MSKCC. Oncoprint analysis (Fig 1D) was performed for CRC AdCA tumors from the PanCan study [73]. Overall survival rates among CRC patients from the TCGA were determined from analysis of microarray data from two cohorts parsed for expression of *PLAGL2*. For one CRC dataset [29] PLAGL2-high tumors were those with greater than 3 SD above the mean for PLAGL2 expression, while another CRC dataset (TCGA-provisional) *PLAGL2*-high tumors were defined as those with greater than 2 SD above the mean for *PLAGL2* expression and *PLAGL2*-low tumors were defined as those with more than 0.25 SD below the mean for *PLAGL2* expression.

### PLAGL2 mutagenesis with CRISPR/eSpCas9 and SRIRACCHA

For expression of eSpCas9 [74], the ORF from eSpCas9 was amplified from eSpCas9(1.1), which was a gift from Feng Zhang (Addgene plasmid # 71814) and cloned into BII-ChBtW, which is identical to BII-ChPtW (which confers Puromycin resistance through PAC), but confers Blasticidin resistance. Using 8 µL Lipofectamine 2000 (ThermoFisher Scientific) this vector, BII-ChBtW-eSpCas9, was introduced into Caco2 and HT29 cells by transfecting 1.6 µg of transposon along with 400 ng of pCMV-hyPBase [75], a generous gift from Dr. Allan Bradley (Wellcome Sanger Institute). Selection with 10 µg/ml Blasticidin was initiated 48 hours after transfection and continued for 7 days, whereupon cells were expanded for transfection with BII-gR-PnGW. The BII-gR-PnGW vector contains a U6-driven gRNA cassette cloned from pX335, which was a gift from Feng Zhang (Addgene plasmid # 42335, [76]). This U6-driven gRNA module was cloned 5’ of the CMV/hEf1a promoter at a unique *Sfi*I site, and was modified to contain two *BsmB*I sites for cloning gRNA oligos, in place of the existing *Bbs*I sites. The BII-gR-PnGW also constitutively expresses GFP-NLS for visualization of transduced cells. Transfections of the BII-gR-PnGW vector were performed in quadruplicate in 24-well plates 24 hours after plating 5 x 10^4^ cells per well. Cells were transfected as above using 400 ng BII-gR-PnGW and 100 ng pCMV-hyPBase, selected with 5 µg/mL Puromycin for 4 days (HT29) or 10 µg/mL Puromycin for 10 days (Caco2). Fluorescence was measured starting 48 hours after transfection. A non-specific gRNA (GGAGACGCTGACCCGTCTCT) was used as a control for comparison with a PLAGL2-targeting gRNA (GTTCACCGCAAGGACCATCTG) [20]. SRIRACCHA-mediated mutagenesis was performed using the BII-gR-PtW-eSpCas9 vector, which contains a gRNA-expression cassette (as in BII-gR-PnGW) and confers resistance to Puromycin. SW480 (8×10^5 cells) or Caco2 (3×10^5 cell) were plated in triplicate or quadruplicate in 6-well plates 24 hours prior to transfection. For one-step transfection with SRIRACCHA components, 600 ng of the BII-gR-PtW-eSpCas9 vector with gRNAs specific for either PLAGL2 (GTTCACCGCAAGGACCATCTG) or LacZ (GAAGGCGGCGGGCCATTACC) were transfected along with 600 ng of the BII-C3H target vector [32] containing target sequences either for PLAGL2 or LacZ cloned at the 3’ end of the PAC ORF, 500 ng of pBS-PtH, and 300 ng of pCMV-hyPBase. SW480 cells were transfected using JetPrime (Polyplus Inc.) (2 µL/µg DNA) per manufacturer instructions while Caco2 cells were transfected using Lipofectamine 2000 (ThermoFisher Inc.) (4 µL/µg DNA) per manufacturer instructions. Cells were selected with Puromycin for 5 days, then switched to Hygromycin selection for 10 days. RNA was then isolated using TRIzol or, alternatively, both RNA and DNA were isolated using the Allprep DNA/RNA Mini Kit (Qiagen Inc.). Mutation analysis was performed either on cDNA (SW480) or gRNA (Caco2). Mutation of *PLAGL2* was determined by amplicon sequencing on the Illumina platform by the Center for Genomic Sciences (Washington University) and analysis was performed using CRISPResso [77]. For transfection with SRIRACCHA components in rescue experiments, Caco2 cells were transfected in a one- or two-step manner. For the two-step method, 3 x 10^5^ cells in a 6-well plate that had been seeded the previous day were transfected with 800 ng BII-C3H-PLAGL2-T1 vector containing the PLAGL2 targeting gRNA sequence, 800 ng of either BII-ChBtW (empty vector), BII-ChBtW-ASCL2 (overexpressing *ASCL2*), or BII-ChBtW-IGF2 (overexpressing *IGF2*), and 400 of pCMV-hyPBase. Transfection medium was removed in 24 hours and replaced with growth medium. Growth medium was changed in a further 24 hours for 10 μg/mL Puromycin and 10 μg/mL Blasticidin containing growth medium, and cells were selected for resistance for 8 days with regular media changes. Once cells reached confluency again, they were trypsinized and removed from the plate, and 3 x 10^5^ cells were again seeded, this time into three identical wells of a 6-well plate. The following day, cells were transfected with 1,600 ng of the BII-gR-PtW-eSpCas9-PLAGL2-T1R vector, the Cas9 activity surrogate vector, 750 ng of the pBS-PtH vector, which contains the hygromycin resistance cassette, and 250 ng of pCMV-hyPBase. Transfection medium was removed and changed as before, and 48 hours post transfection medium containing 400 μg/mL Hygromycin was added. Cells were selected for Hygromycin resistance for 13 days.

For the one-step method, 3 x 10^5^ cells seeded the previous day into a 6-well plate were transfected with 400 ng BII-C3H-PLAGL2-T1 vector containing the PLAGL2 targeting gRNA sequence, 400 ng of either BII-ChBtW (empty vector), BII-ChBtW-ASCL2 (overexpressing *ASCL2*), or BII-ChBtW-IGF2 (overexpressing *IGF2*), 500 ng of the BII-gR-PtW-eSpCas9-PLAGL2-T1R vector, 300 ng of the pBS-PtH vector, and 400 ng of the pCMV-hyPBase vector. Transfection medium was removed and changed again in 24 hours, and 48 hours post transfection cells were selected with 10 μg/mL Puromycin and 10 μg/mL Blasticidin for 5 days, before changing to medium containing 400 μg/mL Hygromycin for 11 days.

For both methods, after Hygromycin selection was finished (cells reached confluency), gDNA was extracted and mutation of *PLAGL2* was again determined by amplicon sequencing on the Illumina platform by the Center for Genomic Sciences (Washington University) and subsequent analysis by CRISPResso.

### Cell proliferation assays

Cellular proliferation was quantified by automated microscopy of GFP-positive cells on a Biotek Cytation3 using a 4x objective, where cell numbers were gauged by enumerating individual cells or by quantifying total integrated fluorescence per well. DLD1 *PLAGL2* mutants [20] express H2BGFP while other cell lines express nuclear GFP following stable transfection with BII-ShPnGW or BII-gR-PnGW. Cellular fluorescence was imaged every 1-2 days and media changed every 2-3 days.

### Cell cycle analysis with FUCCI reporter

A *Piggybac* version of the FUCCI reporter (BII-ChPtW-iresFUCCI) was constructed by PCR amplification of the Clover-Geminin-ires-mKO2-Cdt fragment from pLL3.7 (Addgene Plasmid #83841, [33]) and insertion into unique *BsmB*I sites within the *Piggybac* vector BII-ChPtW. The BII-ChPtW vector was constructed by insertion of the Woodchuck Hepatitis Virus (WHP) Posttranscriptional Regulatory Element (WPRE) downstream of the BsmBI site in the previously described BII-ChPt vector [20]. Transfections of the BII-ChPtW-iresFUCCI vector were performed in quadruplicate in 24-well plates 24 hours after plating 5 x 10^4^ cells per well. Cells were transfected as above using 400 ng of PB transposon and 100 ng pCMV-hyPBase, selected with 5 µg/mL Puromycin for 4 days (HT29) or 10 µg/mL Puromycin for 10 days (Caco2).

### PLAGL2 shRNA-mediated knockdown

For protein knockdown, an shRNA expression *Piggybac* vector, BII-ShPnGW was constructed by insertion of a GFP-NLS cDNA into the BsmBI cloning sites of BII-ChPtW. A U6-driven shRNA module was then inserted at a unique *Sfi*I site upstream of the CMV/hEf1a promoter, with *BsmB*I cloning sites for insertion of unique shRNA double-stranded oligos. The BII-ShPiRW vector was constructed for FUCCI experiments. This vector is identical to BII-ShPnGW, but expresses the iRFP702 near-IR protein [78], which was first codon optimized, synthesized and cloned from a gBlock fragment (IDT Inc.). Alternatively, for rescue experiments (Fig 6), the shRNA module from BII-ShPnGW was cloned as an *Nsi*I-*Age*I fragment into those unique restriction sites in BII-ChPtW-iresFUCCI to generate BII-Sh-FUCCI. The Broad Gene Perturbation Portal was queried for identification of candidate shRNAs against *PLAGL2*, and oligos prepared identical to the strategy employed for shRNA expression from the pLKO.1 vector [79], except overhangs were modified for directional cloning into our BII-ShPnGW vector, with 5’ overhangs ACCG and AAAA at the termini of each dsDNA oligo. The following four *PLAGL2*-specific target sequences were cloned in this manner: shRNA #1: TTCAGGCTCTAGGATCGATTC, shRNA #2: CCGTAGGACTTCAGGTATTAT, shRNA #3: TTGGATGACCTCTAGAGAAAT, shRNA #4: GCAGGAGAGAAGGCCTTTATT. A luciferase-specific shRNA was used as a control, with target sequence TCACAGAATCGTCGTATGCAG. The BII-ShPnGW vector with each specific shRNA was transfected into DLD1 cells in triplicate in 6-well plates, with 1.6 µg of transposon transfected along with 400 ng of pCMV-hyPBase. Selection was initiated 48 hours after transfection using 10 µg/mL Puromycin for Caco2 cells, or 5 µg/mL for other cell lines, and continued for 4-7 days. Cells were then cultured for 1-2 days in medium without Puromycin prior to harvesting RNA or protein for RT-PCR or immunblot.

### Migration and invasion assays

For migration assays 1×10^5^ cells were plated in 0.5 mL DMEM in Falcon Fluoroblok trans-well inserts containing 8 µm pores, which were placed into Falcon 24-well plates, with each well containing 0.75 mL DMEM and 10% FBS. Plates were cultured 24 to 48 hours and then imaged for GFP fluorescence using the Biotek Cytation3. For invasion assays Falcon Fluoroblok trans-well inserts were coated with cold 25% Matrigel (75% DMEM), which was then solidified for one hour at 37°C. Cells were then added to transwell inserts and cultured and imaged as above for migration assays. DLD1 mutant clones (#1 and #2) and parental cells express H2BGFP for visualization [20], while the shRNA knock-down vector (BII-ShPnGW) provided nuclear-localized GFP for enumeration of migratory cells by microscopy using the Biotek Cytation3.

### Soft agar colony formation assays

IEC6 cells were cultured in Advanced-DMEM/F12 (ThermoFisher Inc.) containing 5% FBS, Glutamax (ThermoFisher Inc.), HEPES, N2 Supplement (ThermoFisher Inc.), Plasmocin (Invivogen Inc.), and Penicillin/Streptomycin. Over-expression vectors were prepared by cloning the PLAGL2 coding sequence into BII-ChBtW and IEC6 cells transfected in 6 well plates with 1600 ng of this plasmid and 400 ng of pCMV-hyPBase using Lipofectamine 2000 (ThermoFisher Inc.). Cells were select 48 hours after transfection with medium containing 4 µg/ml Blasticidin and selected for 5 days. To assay anoikis resistance and anchorage-independent proliferation IEC6 cells were grown in 0.35% low-melt agarose (Lonza SeaPlaque). A base layer of agarose was prepared by melting pre-sterilized 1.2% agarose in dH2O at 75°C, and cooling for 60 minutes at 42°C. This was then mixed with pre-heated 0.22 µm-filtered 2x DMEM containing 20% FBS, Sodium Bicarbonate, Sodium Pyruvate, Glutamax (ThermoFisher Inc.), HEPES, 2x N2 Supplement (ThermoFisher Inc.), Plasmocin (Invivogen Inc.), and Penicillin/Streptomycin. This mixture (1 mL) was applied to the bottom of an ultra-low attachment 6-well plate (Greiner Bio-One Inc.) and allowed to cool for 5 minutes at room temperature. Cells were detached with 0.05% Trypsin-EDTA and counted and placed on ice. Pre-sterilized 0.7% agarose was prepared as above, equilibrated for 60 minutes at 42°C, and mixed with pre-warmed growth medium, and then incubated 30 minutes at 37°C. Cells (8×10^4) were added to 3.2 mL of this 0.35% agarose, mixed well, and then 1 mL over-layed onto each well containing a base layer of 0.6% agarose in 6-well plates. Plates were then incubated 20 minutes at room temperature and then placed into a 37°C incubator with 6% CO2. The next day 1 mL of complete culture medium was added. Colonies were counted 3 weeks following plating.

### Organoid culture and transfection

Mouse small intestine organoids (enteroids) were generated from jejunum crypts from three (3) C57BL/6 mice (8-12 weeks) obtained from Jackson Laboratory, and cultured and transfected as previously described [20]. *Plagl2* mutant/KO and PLAGL2 over-expressing (O/E) mouse enteroids were previously described [20]. IGF2 O/E enteroids were generated by cloning the human IGF2 ORF into the BII-ChBtW vector via *Bsm*BI sites, and transfection in mouse enteroids using 800 ng of this BII-ChBtW-IGF2 vector along with 200 ng of pCMV-hyPBase. Enteroid transfections were performed as previously described [20]. Multiple (n=20-50) Blasticidin-resistance clones were obtained and pooled for stable propagation of a polyclonal line. IGF2 expression was assayed by RT-PCR and morphology documented using the Cytation3 microscopy platform and a 4x objective.

### ASCL2 and Wnt reporter assays

The ASCL2 reporter was previously described [38]. Briefly, this reporter was adapted for *Piggybac* transposon-mediated gene delivery by cloning ASCL2 regulatory elements downstream of 2 polyadenylation signals (one synthetic and one from the SV40 TK gene) and upstream of a tandem tomato (tdT) fluorescent protein ORF and NLS signal, followed by the bovine growth hormone (bGH) polyadenylation signal, all flanked by HSIV core insulators from the chicken *HBBA* gene. For transfection, SW480 (2×10^5 cells), HT29 (2×10^5 cells), or Caco2 (1×10^5 cells) were plated in triplicate or quadruplicate in 24-well plates 24 hours prior to transfection. Cells were transfected with 200 ng of BII-STAR-tdT, 200 ng of BII-ShPnGW, and 100 ng of pCMV-hyPBase. The BII-ShPnGW vector contained shRNA sequences directed against a non-specific target or the *PLAGL2* mRNA, as described above. For a positive control of reporter activity BII-STAR-tdT was co-transfected with 200 ng of BII-ChBtW-ASCL2 for ASCL2 O/E. Red fluorescence was quantified 48 hours after transfection using the Cytation3 platform. Cells were then selected with Puromycin for 5 days and then both GFP and RFP fluorescence were quantified using the Cytation3. The BII-TOP-tdT reporter for the canonical Wnt pathway was described previously [20], but was modified for nuclear expression with a C-terminal NLS. This vector was transfected and visualized in SW480, HT29, and Caco2 cells as described above for BII-STAR-tdT. For a positive control of this fluorescent Wnt reporter, BII-ChBtW-dnTCF4 was co-transfected with BII-TOP-tdT. This dominant negative TCF4 variant was described previously [80] and was modified here for expression via transposon. Red fluorescence was quantified 48 hours after transfection using the Cytation3 (Agilent-Biotek Inc.) platform and subsequently after antibiotic selection, as described above for the BII-STAR-tdT reporter.

### FUCCI rescue experiments

For rescuing cell cycle effects caused by PLAGL2 knockdown the FUCCI vector was modified for simultaneous shRNA expression by cloning the *Nsi*I – *Age*I fragment from BII-ShPnGW, which includes shRNA components, into BII-ChPtW-iresFUCCI to create the BII-shFUCCI vector. Caco2 were plated in quadruplicate for transfection with BII-shFUCCI along with BII-ChBtW vectors for O/E of PLAGL2, ASCL2, IGF2, or CTNNB1^S33Y^. The constitutively active beta-catenin mutant (CTNNB1^S33Y^) was described previously [81]. Beginning 48 hours after transfection cells were selected with Puromycin and Blasticidin for 5 days and then RFP and GFP fluorescence were quantified using the Cytation3.

### Immunoblots and protein quantification

For analysis of active and total β-Catenin levels upon *PLAGL2* mRNA KD, Caco2 and SW480 cells in a 6 well dish were transfected with the shRNA construct and selected as previously described, in triplicate. After the initial selection and subsequent expansion of cells to near confluency, cells were harvested and lysate was extracted using Cell Lysis Buffer (Cell Signaling Technology) according to manufacturer’s specifications. Briefly, a 1X concentration of this buffer was made with the addition of 2X protease inhibitor cocktail (Millipore Sigma), 1 mM sodium fluoride, 1 mM sodium pyrophosphate, and 5 mM activated sodium orthovanadate. Cells were rinsed in 1x PBS in the plate, add 500 uL of the lysis buffer was added. The plate was incubated on ice for 5 minutes. The cells and buffer were then removed to a 1.7 mL microcentrifuge tube and sonicated for 10 seconds. Tubes were centrifuged at 14,000 x g for 10 minutes, and supernatant was removed to a separate tube. Protein concentration was determined using a microplate and BCA assay kit (Thermofisher) as per manufacturer’s specifications. Equal amounts of protein (nominally 50 μg) from each sample were separated on 4-20% gradient Bis-Tris SDS-PAGE gels (GenScript), and protein was then electroblotted onto low autofluorescence PVDF membrane (Bio-Rad). The membrane was blocked in 1X PBS + 5% BSA (Millipore Sigma), and probed overnight with primary rabbit antibodies raised against total or active β-Catenin (Cell Signaling Technology, #8480 and #8814 respectively), and primary mouse antibodies raised against α-tubulin (Santa Cruz Biotechnology #sc-5286) as a loading control. The next day, the primary antibodies were washed off and the blot was incubated with goat anti-mouse DyLight 680 conjugated (ThermoFisher #35518) and goat anti-rabbit DyLight 800 (Thermofisher #SA5-10036) conjugated secondary antibodies for an hour. The secondary antibodies were washed off and signal was captured using the LiCor Odyssey CLx Near-Infrared Imaging System, using the 800 channel to capture β-Catenin signal and the 700 channel to capture α-tubulin signal. Quantitation of signal was performed using ImageStudio Lite (LiCor), using the software’s “average” method of background subtraction. β-Catenin signal was normalized using α-tubulin signal, and average normalized β-Catenin signal, standard deviation, and statistical significance were calculated and plotted using GraphPad Prism 8.

## Acknowledgements

We thank the Alvin J. Siteman Cancer Center at Washington University School of Medicine and Barnes-Jewish Hospital in St. Louis, MO. and the Institute of Clinical and Translational Sciences (ICTS) at Washington University in St. Louis, for the use of the Tissue Procurement Core, which provided colon adenocarcinoma and non-malignant tissue samples.

## Supporting information

**Fig S1.**
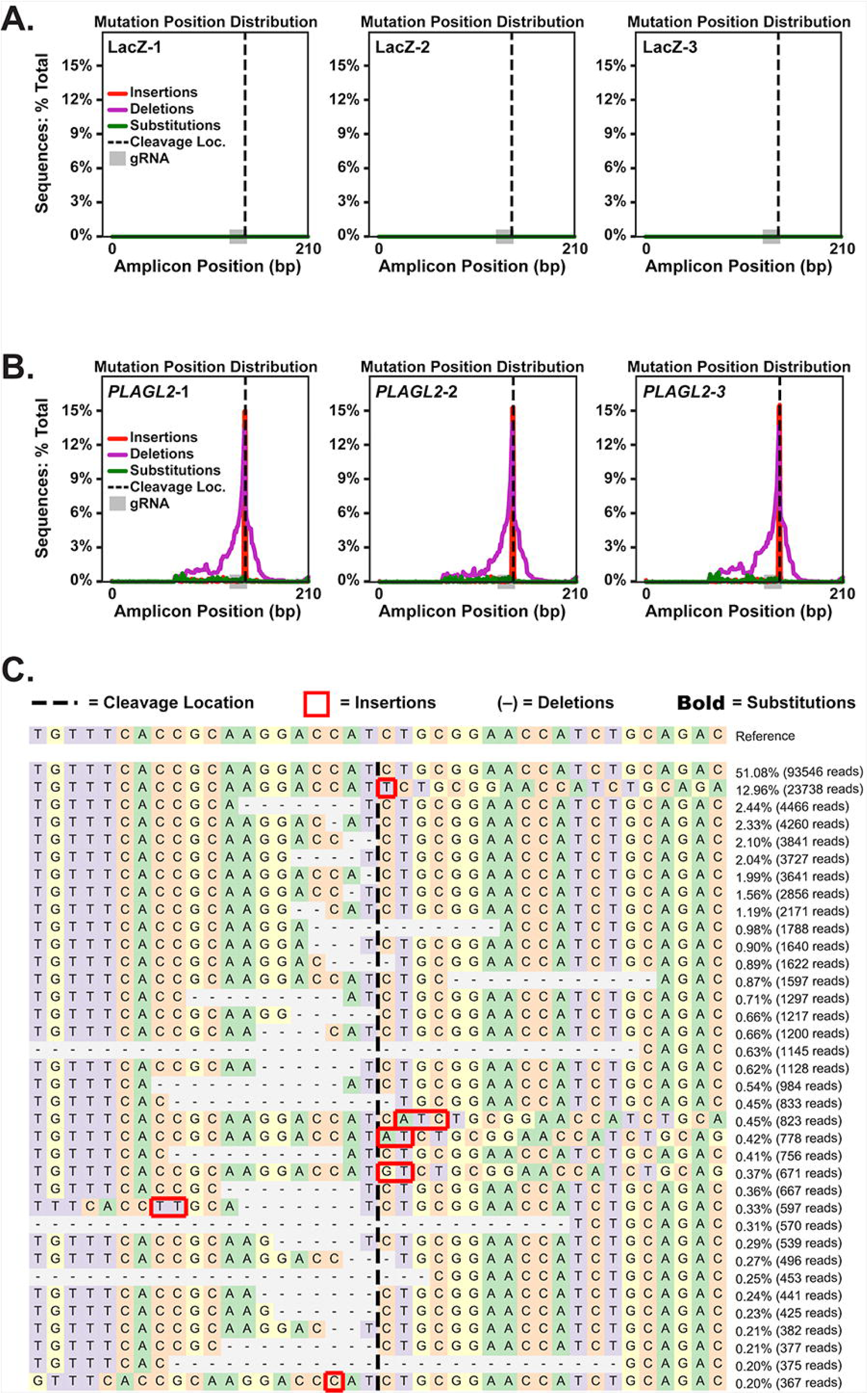
Indel and substitution mutations in SW480 cells around three target SRIRACCHA based CRISPR/Cas9 cleavage sites in *PLAGL2.* **A)** Mutation distribution around cleavage sites in *PLAGL2* gene with NS gRNA, showing no events. **B)** Mutation distribution around the three target cleavage sites in *PLAGL2* **C)** Example allele profile of SRIRACCHA reads from these *PLAGL2* mutations. Note, the top hit represents unmodified *PLAGL2*.

Original Western blot images and raw data used for calculations (means, statistics, etc.) can be found at https://wustl.box.com/s/288v1qk8ksiqxl4wyufy44mtq4tj2c3p.

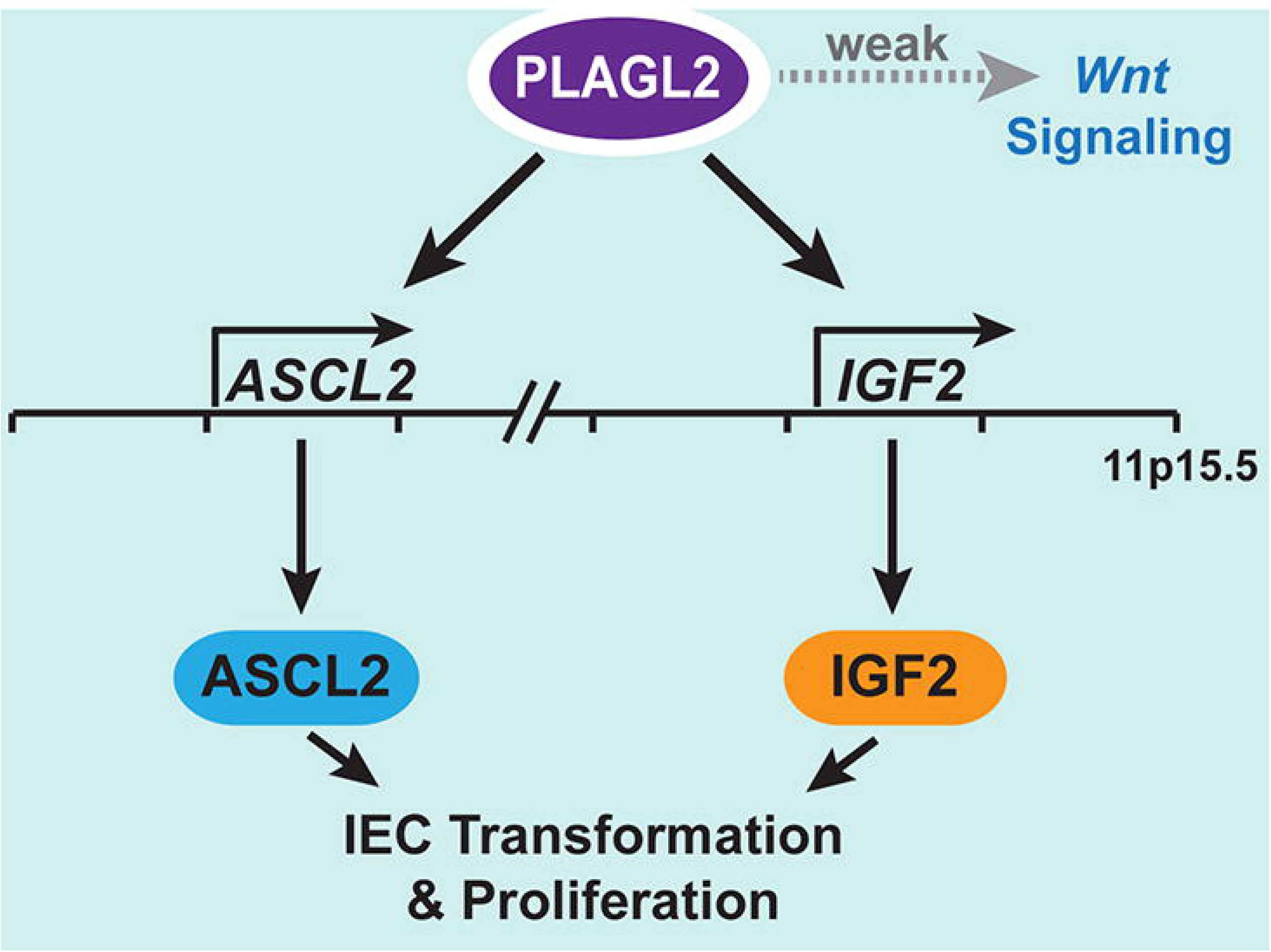

